# To explore the role of NABP2 in the immune microenvironment and metabolic pathway of liver cancer

**DOI:** 10.64898/2025.11.30.691448

**Authors:** Ying Li, Yibing Han, Linyi Cheng, Jiayi Zhang, Ruolan Deng, Wen Qiu

## Abstract

**Objective:** This study aims to explore the specific signaling pathways associated with the NABP2 gene and its potential molecular mechanisms influencing the progression of hepatocellular carcinoma (HCC), thereby identifying new therapeutic targets.

**Method:** We collected 50 normal and 374 tumor samples to investigate the differential expression of NABP2 in HCC. Variousgenomic, single-cell, and bioinformatic analyses, including gene microarray technology, were employed. Data processing and analysis were conducted using R language, focusing on the correlation between NABP2 expression, immune cell infiltration, drug sensitivity, and metabolic pathways.

**Results:** Our findings indicate that NABP2 plays a significant role in the development of HCC and the tumor immune microenvironment. Analyses such as GSVA and GSEA suggest that NABP2 may influence disease progression through involvement in key molecular signaling pathways, including DNA replication, mRNA splicing, and protein degradation. Furthermore, a notable **correlation** was observed between NABP2 and various metabolic pathways, as well as specific immune cell populations. Additionally, we identified potential chemotherapeutic agents for HCC based on drug sensitivity analysis.

**Conclusion:** These findings position NABP2 as a crucial biomarker for the progression and treatment of hepatocellular carcinoma, offering new insights into the pathogenesis and prognostic strategies for HCC in future research.

---

Currently, cancer has become one of the leading causes of death worldwide, among which the mortality rate of Hepatocellular carcinoma ranks fourth [1]. According to the Chinese Guidelines for the Integrative Diagnosis and Treatment of Tumors (CACA) - Hepatocellular carcinoma section, the incidence rate of Hepatocellular carcinoma in China was as high as 18.2 per 100,000 population in 2018 [2], second only to lung cancer, stomach cancer, and breast cancer, ranking fourth. Clinically, Hepatocellular carcinoma is divided into primary Hepatocellular carcinoma, secondary Hepatocellular carcinoma, and other malignant tumors of the liver. Among them, primary Hepatocellular carcinoma is the most common, and Hepatocellular carcinoma accounts for a very high proportion of primary Hepatocellular carcinoma [3]. Hepatocellular carcinoma (HCC), originating from liver cells, is a malignant tumor that usually forms one or more tumors in liver tissue. Current relevant data indicate that the main pathogenic cause of Hepatocellular carcinoma is hepatitis virus infection, and other pathogenic factors include drinking polluted water sources, consuming food contaminated with aflatoxin, liver cirrhosis, fatty liver, non-alcoholic fatty liver disease [4], genetic factors, etc. [5] According to epidemiological surveys, the male-to-female prevalence ratio ranges from 2:1 to 4:1, indicating that the prevalence rate in females is significantly lower than that in males [6]. It is worth noting that in the early stage of Hepatocellular carcinoma, patients usually have no obvious symptoms. As the disease progresses, symptoms such as loss of appetite, weight loss, persistent pain in the liver area, hepatomegaly, persistent fever, and jaundice begin to appear. Common diagnostic methods for Hepatocellular carcinoma include liver Function Tests, liver ultrasound, CT scans, MRI, andliver biopsy. The key to preventing Hepatocellular carcinoma is to avoid high-risk factors, such as maintaining good liver health, receiving hepatitis B vaccination, limiting alcohol intake, maintaining a healthy weight and diet, and avoiding the use of unregulated needles [7]. Regular screening and diagnosis for high-risk groups of Hepatocellular carcinoma play an important role in prompting the timely detection of Hepatocellular carcinoma. Currently, the most convincing early screening methods are liver ultrasound (alpha-fetoprotein, AFP) and serum alpha-fetoprotein levels. Additionally, upper abdominal computed tomography, liver function tests, and viral screening can be performed, and liver biopsy may be conducted if necessary [8]. Therefore, it is recommended that high-risk individuals undergo examinations every 6 months.

It is worth noting that maintaining genomic stability is crucial for preventing Cancer development. As a member of the single-stranded DNA binding protein (SSB) family, NABP2 (nucleic acid binding protein 2, also known as hSSB1, OBFC2B, SOSS-S1) plays a central role in DNA replication, DNA damage repair, and maintenance of genomic stability [9] [10] [11]. Given its critical role in genomic integrity, investigating the specific molecular mechanisms of NABP2 in the occurrence and development of Liver cancer, particularly its association with tumor Immune microenvironment remodeling and metabolic reprogramming, is of great significance for identifying new therapeutic targets.

Gene chip, also known as DNA microarray technology, is a mature and widely used biological detection technology [12]. In recent years, bioinformatics analysis has been widely applied to microarray data to identify key genes and conduct subsequent analyses. Gene chip technology is a high-throughput DNA detection technology based on the principle of nucleic acid complementarity, which takes gene sequences as the analysis object and is commonly used for large-scale screening of differentially expressed genes [13]. In this study, Gene chip technology was used to analyze the co-expression of NABP2 gene in Liver cancer samples and its differential expression levels between normal tissues and cancer tissues. Based on gene chip technology, we identified the NABP2 gene and further explored and analyzed the specific Signaling pathways involved and its potential role in influencing tumor progression molecular mechanisms, aiming to find new breakthroughs for the treatment of liver cancer.

## 1 Materials and Methods

### 1.1 Co-expression Analysis

This study simultaneously analyzed the co-expression of the NABP2 gene in Liver cancer data, with the correlation coefficient set to 0.4 to exclude other data and the p-value set to 0.05. genes significantly co-expressed with NABP2 were screened, their correlations were analyzed, and circle plots and heatmaps were generated using the "Corrplot" and "Circlize" packages.

### 1.2 Immune cell infiltration Analysis

CIBERSORT is a computational method used to evaluate the relative abundance of different cell types in complex mixed tissue samples. Unlike other computational methods, it applies deconvolution analysis to the expression matrix of immune cell subgroups based on the principle of support vector regression. It can distinguish human immune cell phenotypes such as plasma cells and T cells through 547 biomarkers.cells, and myeloid cell subsets, totaling 22 types [14]. In this study, the CIBERSORT algorithm was used to analyze RNA-seq data from Liver cancer patients, The relative proportions of 22 types of human immune infiltrating cells were calculated, and the correlation between the content of various immune cells and the expression levels of related genes was analyzed. It was determined that a statistical significance exists when P < 0.05.

### 1.3 gene Set Variation Analysis (GSVA)

Gene set variation analysis (GSVA) performs unsupervised classification of samples and then transforms the expression matrix of target genes among different samples into the expression matrix of gene sets among samples, taking gene sets as the basic unit of analysis and using non-parametric statistical methods to explore changes in biological pathways or biological functions [15]. Therefore, we downloaded gene sets from the MSigDB database and used the GSVA algorithm to conduct comprehensive scoring, and evaluated the potential biological function changes of gene sets in different samples through the final scores.

### 1.4 gene Set Enrichment Analysis (GSEA)

GSEA can deeply analyze the differences in Signaling pathways between lowly expressed genes and highly expressed genes. Therefore, in this study, patients were divided into high and low expression groups based on their NABP2 expression levels to investigate pathway differences [16]. For reference genes analyzing differential pathway expression between subgroups, we selected the fully annotated gene Set Version 7.0 downloaded from the MsigDB database. Based on consistency scores, significantly enriched gene sets were ranked, with the p-value adjusted to less than 0.05. GSEA (Gene Set Enrichment Analysis), as a commonly used method, was employed to conduct in-depth research on the association between disease subtypes and biological significance.

### 1.5 Drug sensitivity Analysis

The Genomics of Drug sensitivity in Cancer (GDSC) database (https://www.Cancerrxgene.org/) collects the sensitivity and response of various tumor cells to drugs, serving as the largest public resource for tumor cell Drug sensitivity to date. In this study, the R package "oncoPredict" and regression analysis methods were used to predict the chemotherapy sensitivity of each tumor sample and the IC50 value for each specific chemotherapy drug treatment. Additionally, 10-fold cross-validation was performed on the GDSC training set to verify the correctness of the regression analysis and the accuracy of the predicted values. Throughout the process, defaults such as "combat" for removing batch effects and the mean value for duplicate gene expressions were applied using the average.

### 1.6 Analysis of TMB, MSI, and NEO Data

TMB refers to the total number of mutations present in a tumor, including detected somatic gene coding errors, base insertions, deletions, or substitutions. In this study, TMB was defined by calculating the mutation frequency and number of mutations/exon length for each tumor sample, and dividing the number of non-synonymous mutation sites by the total length of the protein-coding region. The MSI values for each TCGA patient were derived from a previously published study [17]. NetMHNABP2an v3.0 was used to evaluate the Neoantigen of each patient [18].

### 1.7 Construction of Nomogram Model

Nomogram is based on multivariate regression analysis, which integrates clinical symptoms of the disease and the measured content of gene expression into scaled line segments and draws them on the same plane according to a scale, thereby expressing the mutual relationship between variables in the prediction model.as described in [19]. In this study, each influencing factor was assigned a score based on different contribution degrees (the magnitude of regression coefficients), the degree of influence of each factor on the outcome variable. The total score was then obtained by summing up the individual scores, and the predicted value was subsequently calculated.

### 1.8 Single-cell data quality control

First, we used the "Seurat" package to read in the expression profiles, and then performed cell filtering based on the number of expressed genes per cell, total UMI count, and mitochondrial expression proportion per cell. The mitochondrial expression proportion of a cell refers to the percentage of the expression level of mitochondrial genes in a cell relative to the total expression level of all genes. When a cell begins to enter the death program, it indicates a high proportion of mitochondria and a low proportion of RNA in its gene expression. We performed quality control using the median absolute deviation (MAD), and it is generally considered that if a variable is more than 3 MAD away from the median, the value is regarded as an outlier and needs to be removed subsequently, we used DoubletFinder (V2.0.4) to filter out doublets from each sample respectively, and thus completed cell quality control [20].

### 1.9 Dimensionality reduction and clustering of single-cell data

First, this study applied the global normalization method LogNormalize to standardize the single-cell data, which involves multiplying the data by a coefficient s0, then adjusting the total expression of each cell to 10000, and finally taking the logarithm of the total expression; CycleScoring of cell was used to calculate cell cycle scores; FindVariableFeatures was employed to identify highly variable genes; furthermore, since the proportion of mitochondrial gene expression and UMI counts caused fluctuations in gene expression, we selected the ScaleData function to eliminate such fluctuations; RunPCA was applied to perform linear dimensionality reduction on the expression matrix, and the main components were selected for subsequent analysis; Harmony was used to remove batch effects, and RunUMAP was employed for uniform manifold approximation and projection (UMAP) for nonlinear dimensionality reduction [21].

### 1.10 cell annotation

Cell annotation was performed by identifying the cell types present in the corresponding tissues and their corresponding marker genes, primarily through querying the CellMarker and PanglaoDB databases and literature, supplemented by automated annotation using SingleR software. We used the FindAllMarkers function to screen marker genes for each category. The screening parameters were: only.pos = TRUE, min.pct = 0.25, logfc.threshold = 0.25. That is, only positive marker genes were screened, and the marker genes were expressed in at least 25% of the cells, with a LogFC threshold of 0.25 for differential expression.

### 1.11 Ligand-receptor interaction analysis (CellChat)

cellChat is a tool for analyzing intercellular communication in single-cell RNA sequencing data, which infers intercellular communication by simulating the probability of cell-cell communication and combining gene expression data with the relationships between signaling ligands, receptors, and their cofactors signaling network [22]. In this analysis, the study used the normalized single-cell expression profiles as input data and the cell subgroups derived from single-cell analysis as cell information. It analyzed cell-related interactions and quantified the closeness of interaction relationships using intercellular interaction strength (weights) and frequency (count), thereby observing the activity level and influence of each cell type in the disease.

### 1.12 Statistical analysis

All statistical analyses mentioned above were performed using R language (version 4.3.0), with statistical significance set at p<0.05.

## 2 Results

### 2.1 TCGA Data Acquisition and Differential Analysis

The TCGA database (https://portal.gdc.cancer.gov/) is a large-scale cancer gene information database jointly created and maintained by NCI and NHGRI, which collects a large number of cancer samples, performs multi-dimensional analyses such as genome, transcriptome, and epigenome through high-throughput sequencing technology, and stores various genetic information data including SNP data, gene expression data, and copy number variations. We downloaded the processed original mRNA expression data of liver cancer from the database. A total of 424 manifestations and rootcauses were collected, including normal samples (n=50) and tumor samples (n=374) (Figure 1), which were used for the differential analysis of NABP2 expression. The expression of this gene was visualized and analyzed using the GSE14520 (GPL3921, 220 vs 225) dataset from the GEO database, and the results were consistent with the analysis results for visualization analysis, and the results were consistent with the analysis results.

**Figure 1.**
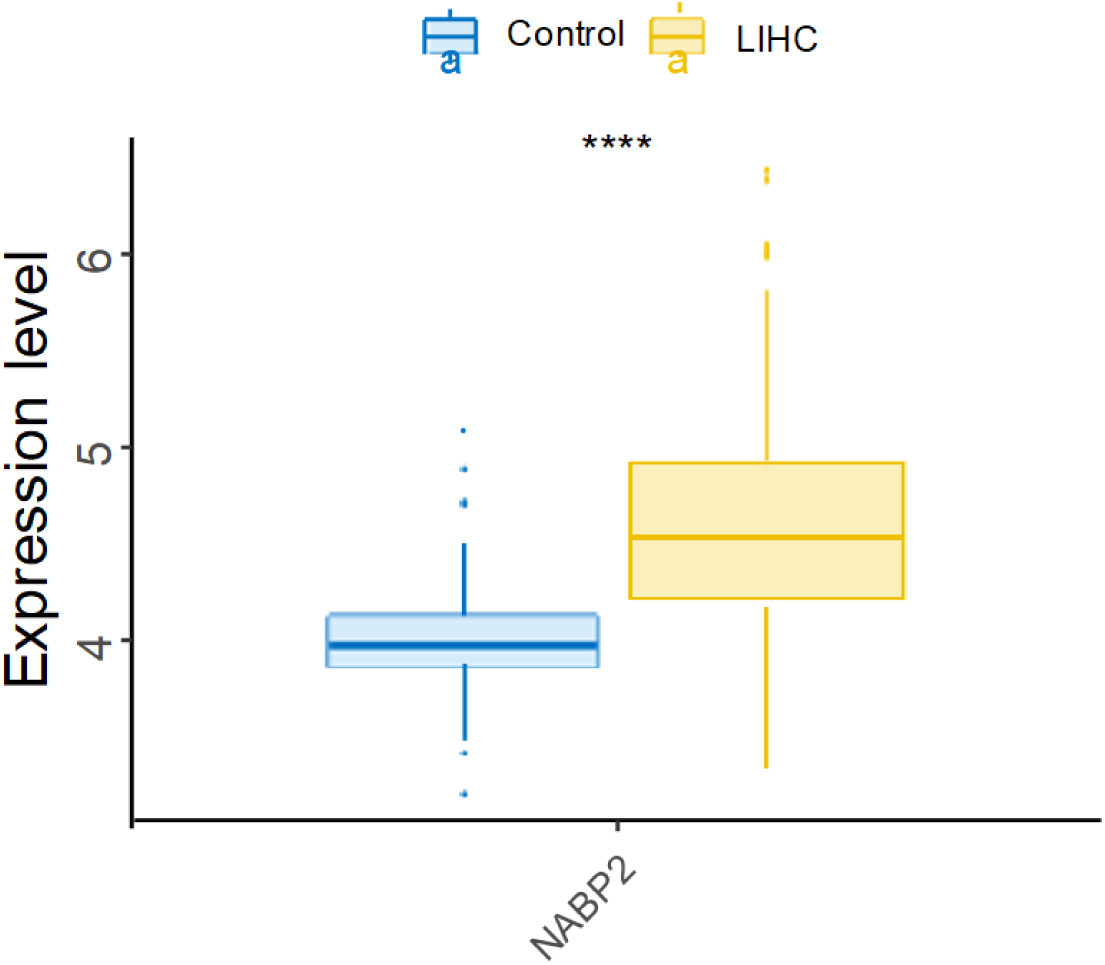
Supplementary public dataset

The GEO database (https://www.ncbi.nlm.nih.gov/geo/info/datasets.html), whose full name is GENE EXPRESSION OMNIBUS, is an international public repository created and maintained by NCBI that contains various high-throughput genomic data including gene expression profiles and non-coding RNA analysis. Single-cell data files of GSE202642 were downloaded from the NCBI GEO public database, and expression profile data from a total of 11 patients were included.

### 2.2 The specific mechanism of NABP2 in liver cancer

We downloaded the original mRNA expression data of liver cancer from the TCGA database. Survival analysis grouped by high and low expression levels of the NABP2 gene showed that the P value of NABP2 was (p = 0.0001) when taking the median survival time (Figure 2A). Next, differential expression level analysis of NABP2 revealed a significant difference in NABP2 expression between normal tissues and cancer tissues, with a significant increase in tumor samples (Figure 2B). We established univariate and multivariate Cox regression models based on NABP2 expression and various clinical data and plotted forest plots (Figure 3). In addition, we analyzed the relationship between NABP2 and clinical symptoms, and the results showed that NABP2 was associated with Fustat, Grade, Stage of patients, and T stage was statistically significant.

**Figure 2.**
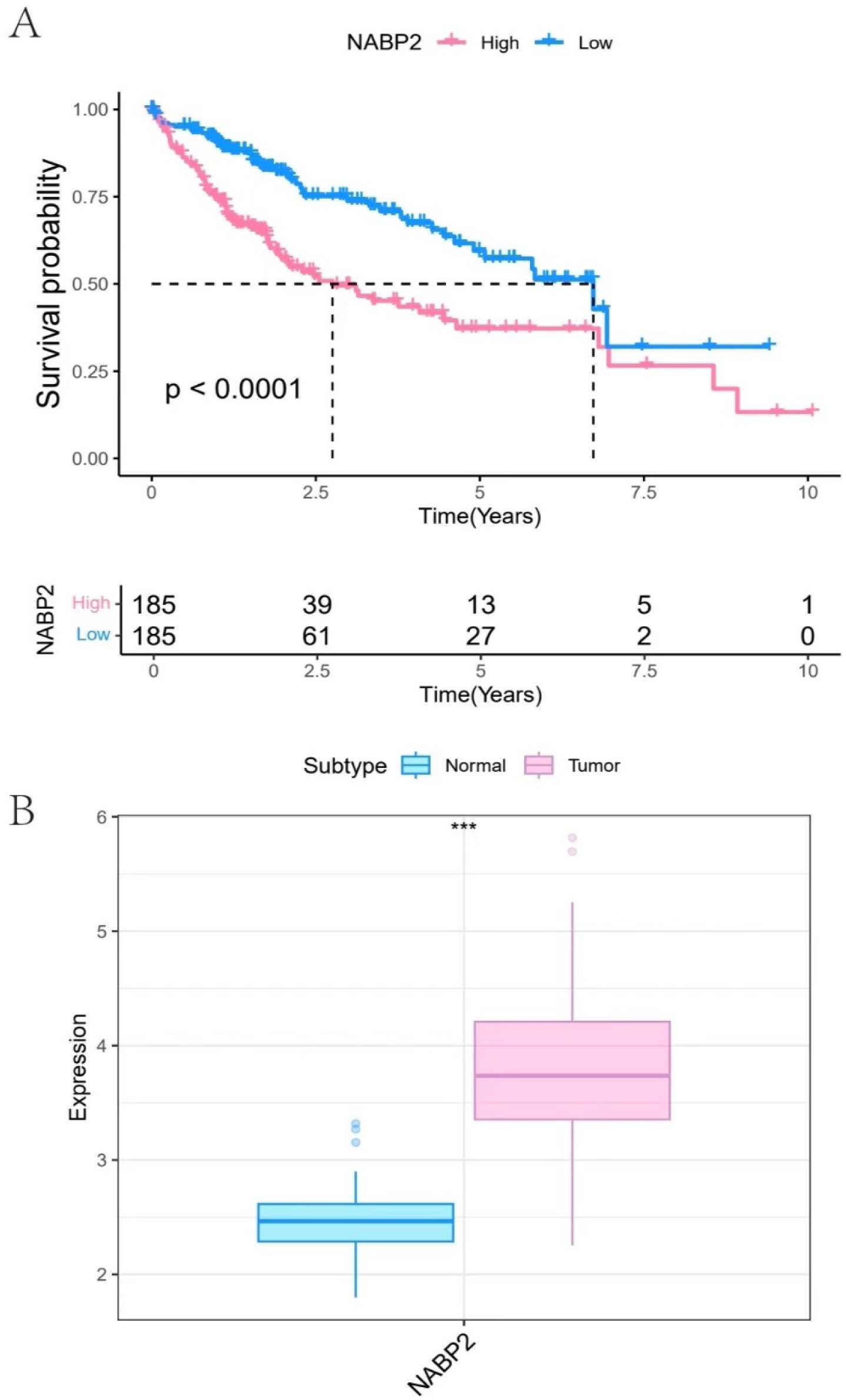
Survival analysis and expression difference of NABP2 gene (A) Survival status of NABP2 gene. (B) Expression difference of NABP2 gene, where blue represents control patients and pink represents tumor patients.

**Figure 3.**
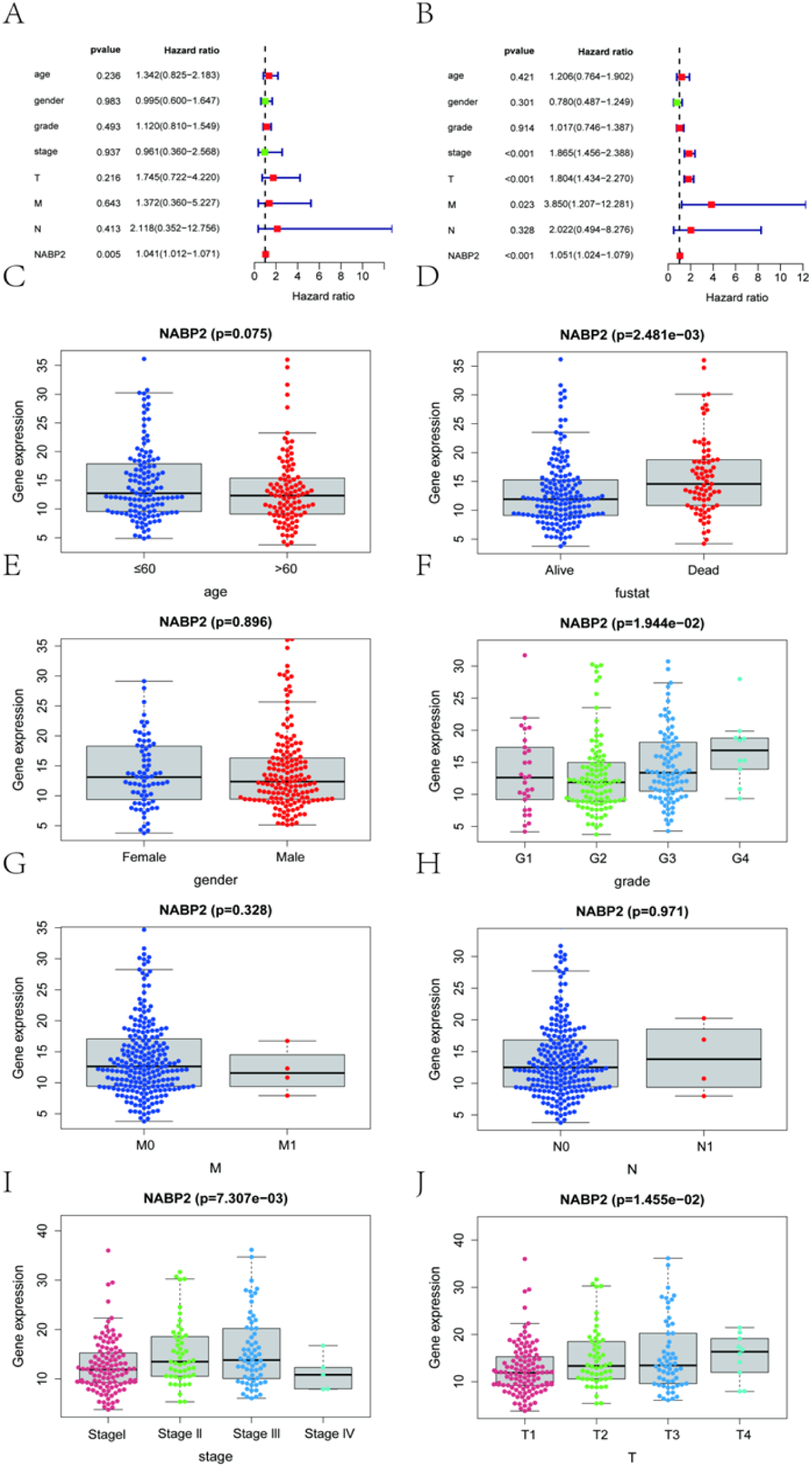
Univariate and multivariate clinical correlations of NABP2 gene (A-B) In the univariate Cox regression analysis, NABP2 was a risk factor in liver cancer patients and was statistically significant. (C-J) Correlation between NABP2 gene and clinical characteristics

### 2.3 Co-expression of the NABP2 gene

Based on the expression profiles of liver cancer patients in The Cancer Genome Atlas (TCGA) database, we further explored the co-expression network of NABP2 through correlation analysis, with a correlation coefficient filtering threshold of 0.4 and a p-value of 0.05. A total of 2285 genes significantly correlated with NABP2 expression were selected. A heatmap of the top 10 genes with positive/negative correlation coefficients (Figure 4A) and a circle diagram of gene co-expression correlations (Figure 4B) were generated.

**Figure 4.**
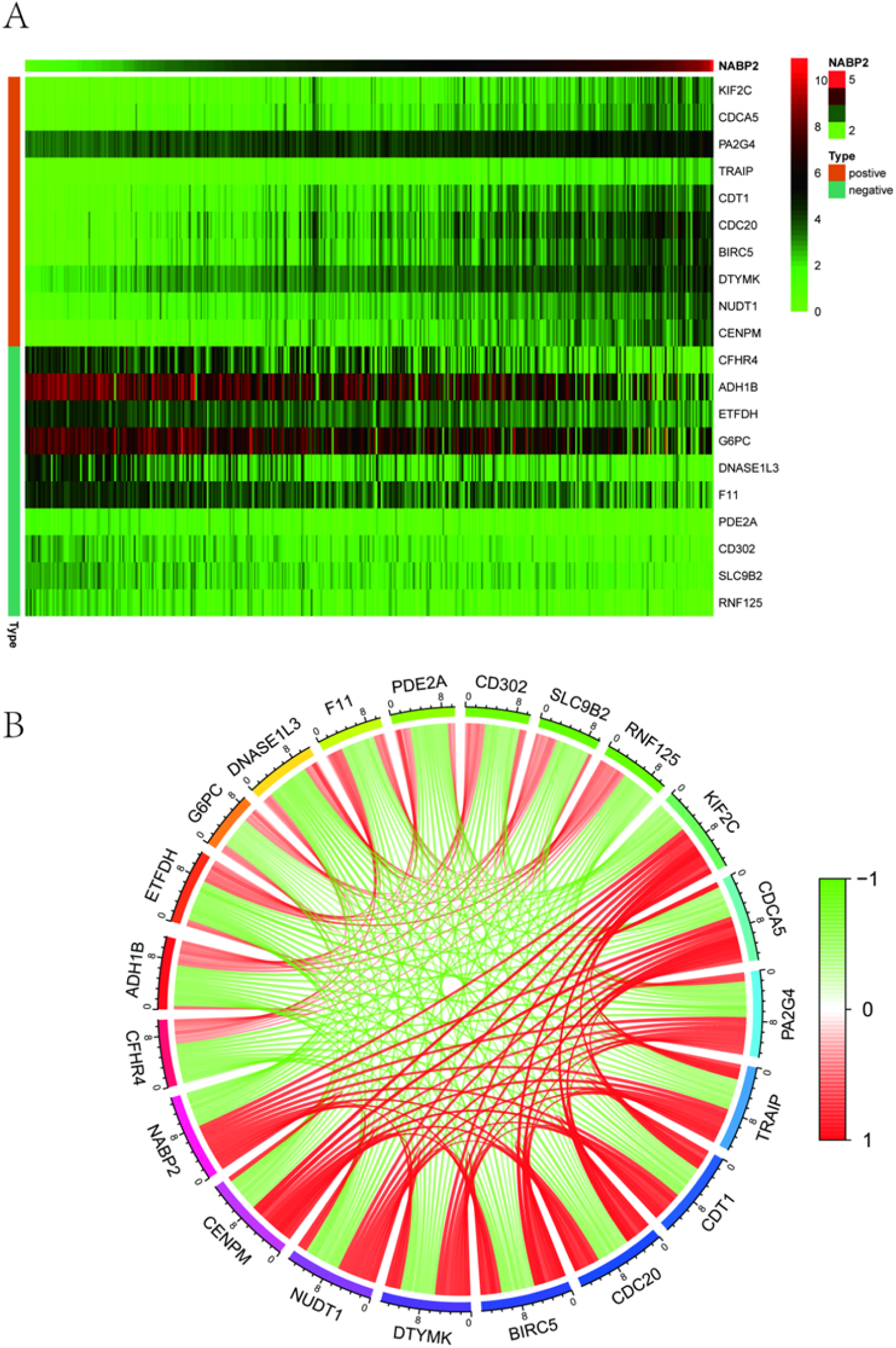
Co-expression correlation analysis of NABP2 (A) Heatmap of the top 10 genes with positive/negative correlation coefficients. (B) Co-expression correlation circle plot of the top 10 genes with positive/negative correlation coefficients, where red represents positive correlation and green represents negative correlation.

### 2.4 Relationship between NABP2 and Tumor Immune Infiltration

The tumor microenvironment includes the internal and external environment where tumor cells reside, as well as the structure, function, and metabolism of the tissue where they are located, namely the extracellular matrix, immune cells, growth factors, inflammatory factors, and tumor cells themselves. The tumor microenvironment plays a crucial role in tumor diagnosis, assessment of patient survival status, and clinical treatment sensitivity [23]. By analyzing the relationship between NABP2 and tumor immune infiltration in the TCGA dataset, this study demonstrated the correlations between immune cells and the proportion of immune cells in each patient in various forms. (Figure 5A-B) We find that B cells memory、B cells naive、Dendritic cells resting、Eosinophils、Macrophages M0、Macrophages M2、 Mast cells activated、Monocytes、NK cells resting、Plasma cells、T cells CD4 memory activated、T cells CD4 memory resting、T cells follicular helper、T cells regulatory (Tregs) have significant meanings between the two groups at the same time (Figure 5C). It is worth noting that high expression of NABP2 is significantly associated with increased infiltration of immunosuppressive cells (such as Tregs and M2 macrophages), while negatively correlated with some cells with anti-tumor potential (such as activated CD4+ T cells), suggesting that NABP2 may be involved in shaping the immunosuppressive microenvironment. The relationship between the NABP2 gene and immune cells will be the direction of our next research(Figure 5D).

**Figure 5.**
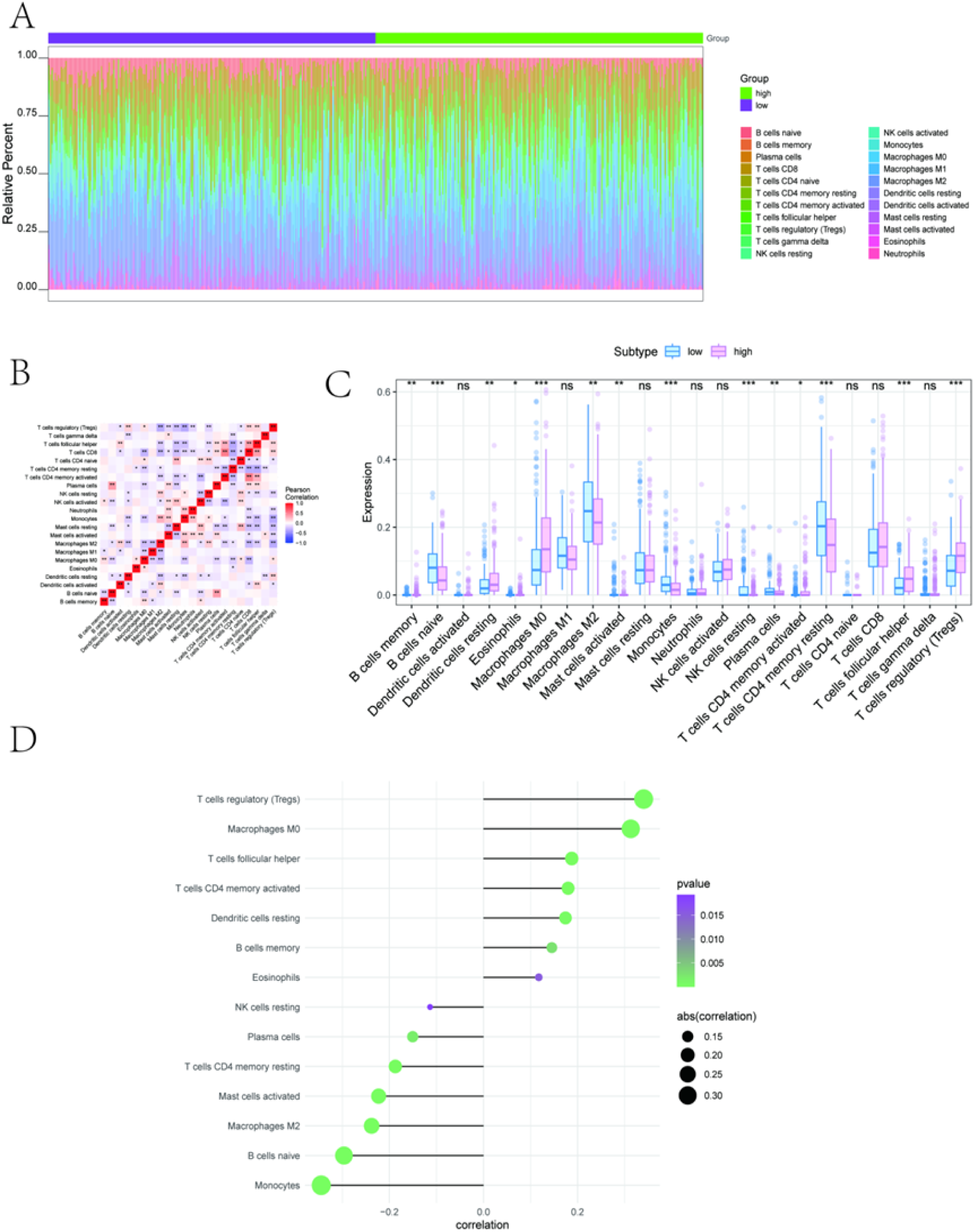
Relationship between NABP2 and tumor immune infiltration (A-B) Bar charts of relative percentages of immune cells and correlation heatmaps. (C) Expression differences of immune cell subgroups in high/low groups. (D) Correlation analysis between NABP2 and immune cells

### 2.5 Potential molecular mechanism of NABP2 affecting tumor progression

Based on the signaling pathways involved in the NABP2 gene discovered in the study, we further explored the potential molecular mechanism by which the NABP2 gene affects tumor progression. GSVA results showed that patients with high NABP2 expression could enrich signaling pathways such as E2F_TARGETS, DNA_REPAIR, and MYC_Targets_V1. In addition, GSEA results indicated that NABP2 could enrich signaling pathways including DNA replication, Proteasome, and Spliceosome (Fig. 6A-B). The charts showed that NABP2 might affect liver cancer progression through these pathways. We downloaded the processed SNP-related data of liver cancer, selected the TOP30 genes with higher mutation frequencies for display, compared the differences in mutated genes between the two groups of patients, and plotted the mutation landscape using the R package ComplexHeatmap (Fig. 6C). The image indicated that genetic mutations such as TP53 in patients with the high expression group. The proportion was significantly higher in the high-expression group compared to the low-expression group.

**Figure 6.**
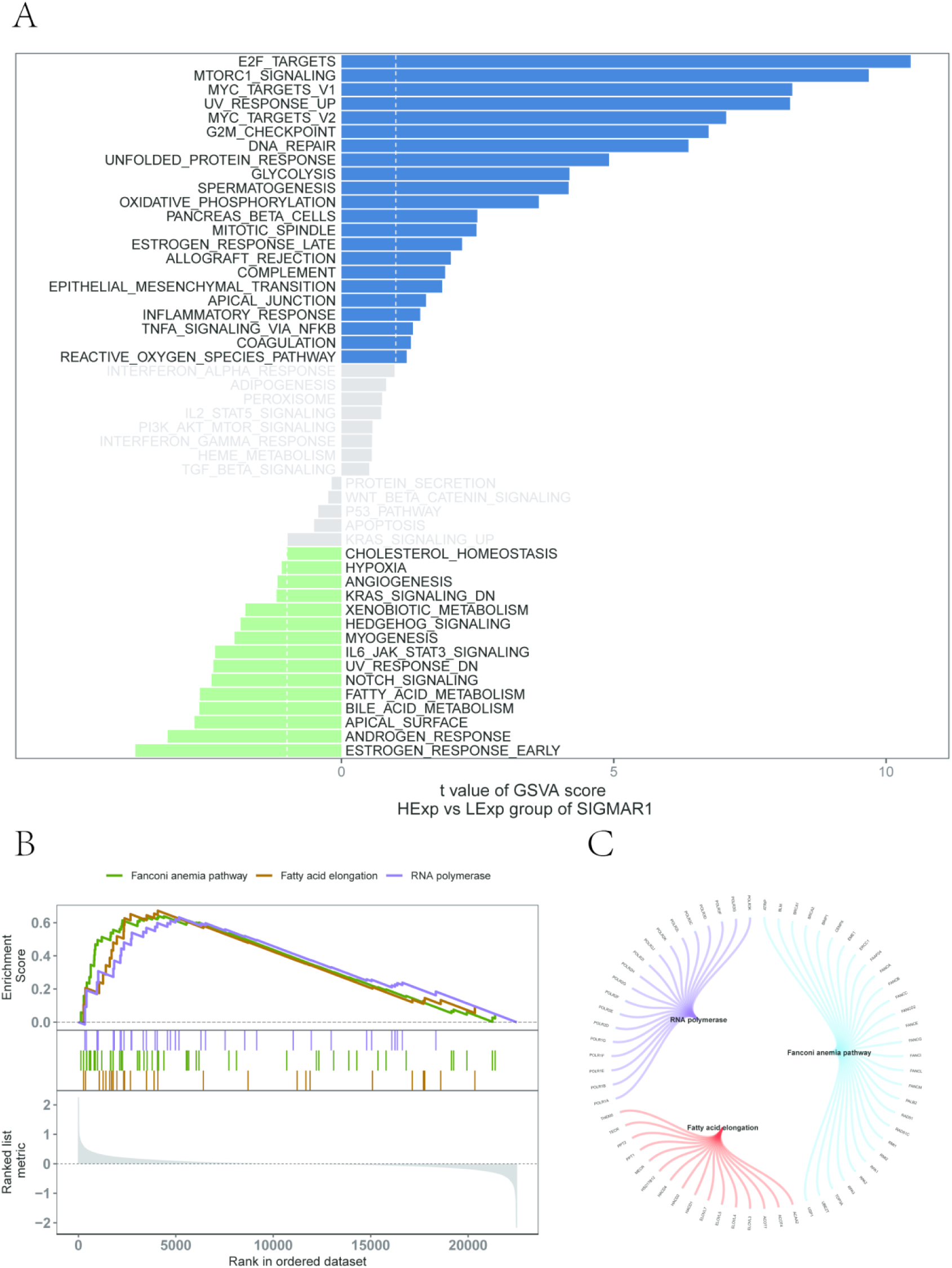
GSVA and GSEA analysis (A) The signaling pathways involved in the NABP2 gene, with green indicating the signaling pathways involved in low-expression genes, and the background gene set is hallmark. (B-C) The KEGG signaling pathways involved in the NABP2 gene, as well as pathway regulation and the genes involved.

### 2.6 NABP2 and drug therapy

The method of surgery combined with chemotherapy has a good effect on early-stage liver cancer. We used "oncoPredict" to predict the chemosensitivity of each tumor sample, and these drug sensitivity data were all from the GDSC database to further explore the relationship between NABP2 and the sensitivity of common chemotherapeutic drugs. According to the drug sensitivity analysis (all p < 0.05), the expression of NABP2 was significantly positively correlated with the sensitivity of Camptothecin-1003 and Vinblastine-1004 (i.e., high expression of NABP2,NABP2 expression was significantly positively correlated with the sensitivity to Camptothecin-1003 and Vinblastine-1004 (i.e., high NABP2 expression was associated with high IC50 values and low sensitivity), and significantly negatively correlated with the sensitivity to Cisplatin-1005, Cytarabine-1006, Docetaxel-1007, and Gefitinib-1010 (i.e., high NABP2 expression was associated with low IC50 values and high sensitivity) (Figures 7A-B). We further explored the relationship between the NABP2 gene and common immunotherapy-related tumor markers and found significant differences in microsatellite instability (MSI) and tumor mutation burden (TMB) between the high and low NABP2 expression groups (Figure 7C).

**Figure 7.**
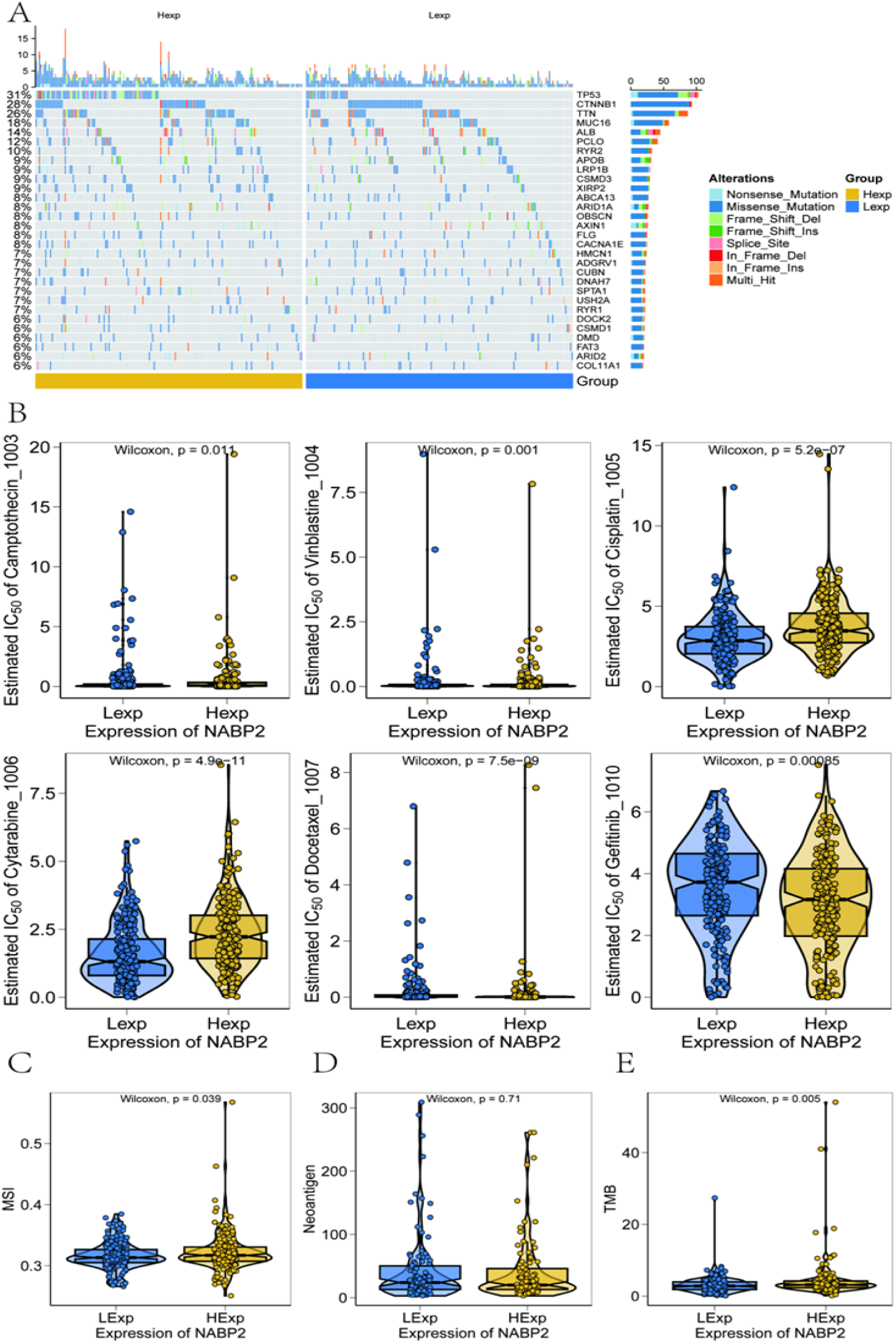
Correlation between tumor mutations and drug sensitivity (A) SNP-related data of liver cancer, displaying the TOP30 genes with higher mutation frequencies to compare the differences in mutated genes between the two patient groups. (B) Analysis of the sensitivity between NABP2 gene and chemotherapeutic drugs. (C-E) Differences in microsatellite instability (MSI), tumor mutation burden (TMB), and tumor neoantigen between high and low NABP2 expression groups.

### 2.7 NABP2 Expression and Model Prediction

We presented the results of their regression analysis in the form of a nomogram based on NABP2 expression levels. The regression analysis results indicated that in all our samples, the values of different clinical indicators of liver cancer and the distribution of NABP2 expression contributed to varying degrees throughout the scoring process (Figure 8A). Meanwhile, we also conducted predictive analysis on the overall survival (OS) for two periods, one year and three years (Figure 8B). The results showed that the predicted OS was in good agreement with the observed OS, indicating that the Nomogram model has good predictive performance.

**Figure 8.**
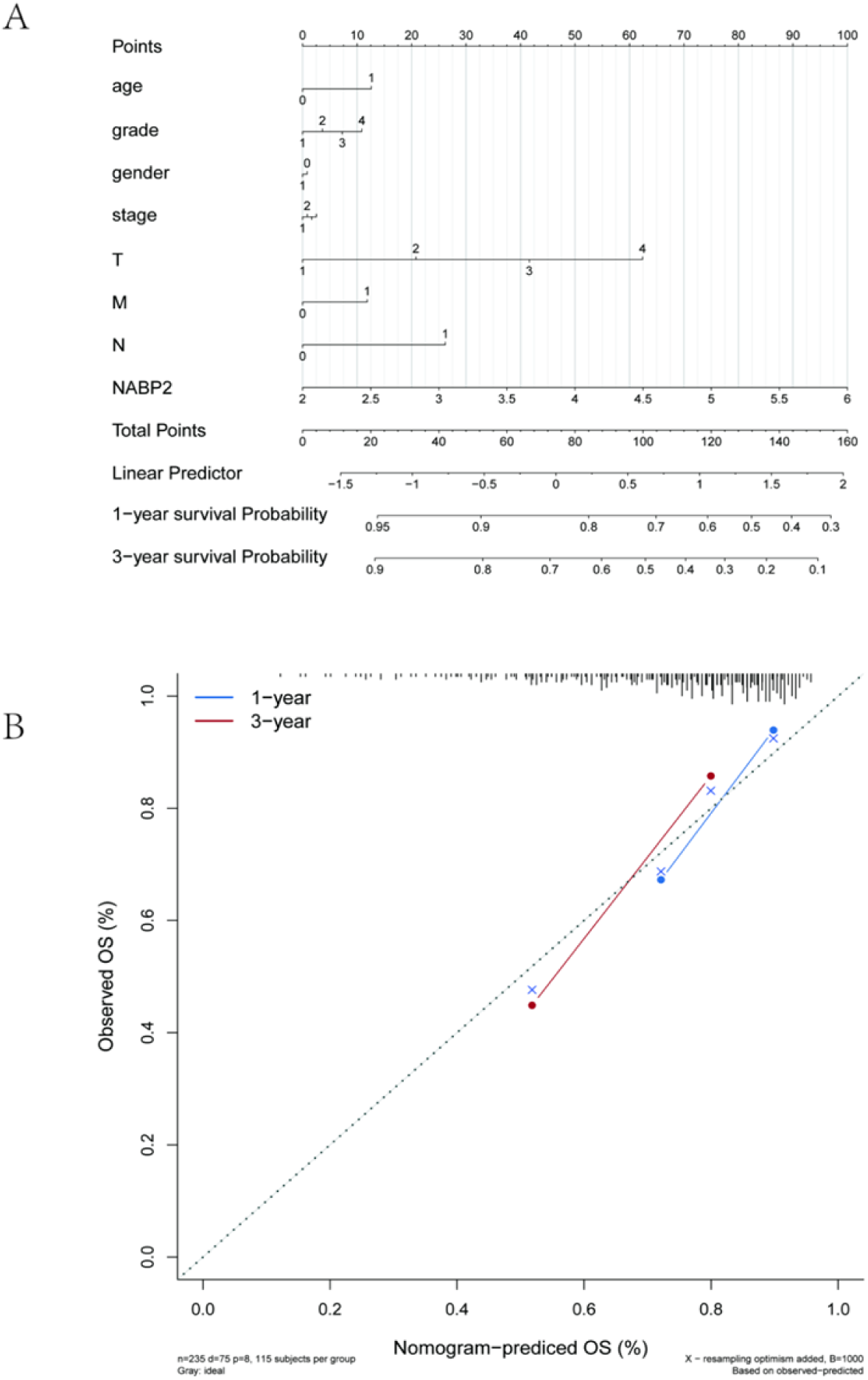
Construction of the Nomogram model (A) The values of different clinical indicators for liver cancer and the expression distribution of NABP2 contribute to varying degrees throughout the scoring process. (B) Predictive analysis of OS for the 1-year and 3-year periods.

### 2.8 Acquisition of 10 cell subsets

Single-cell data of GSE202642 were downloaded from the NCBI GEO public database, with a total of 11 samples included.

nFeature_RNA and nCount_RNA were used for preliminary screening of data samples (nFeature_RNA >= 200 & percent.mt <= median+3MAD & nFeature_RNA <= median+3MAD & nCount_RNA <= median+3MAD). Subsequently, 2000 highly variable genes were screened out through analysis of variance, with a standard deviation range of [0.8, 15.2], and the top 10 genes with the highest standard deviations were displayed (Figure 9A). The data were sequentially subjected to standardization, normalization, PCA, and Harmony analysis (Fig. 9B-D). PCA analysis showed that the first 30 principal components cumulatively explained 85% of the variance. The Harmony algorithm successfully integrated multi-sample data, reducing the batch mixing index from 0.68 to 0.12. Finally, after UMAP nonlinear dimensionality reduction analysis, cells formed 12 topologically separated subpopulations in two-dimensional space, indicating significant transcriptional heterogeneity (Fig. 9E).

**Figure 9.**
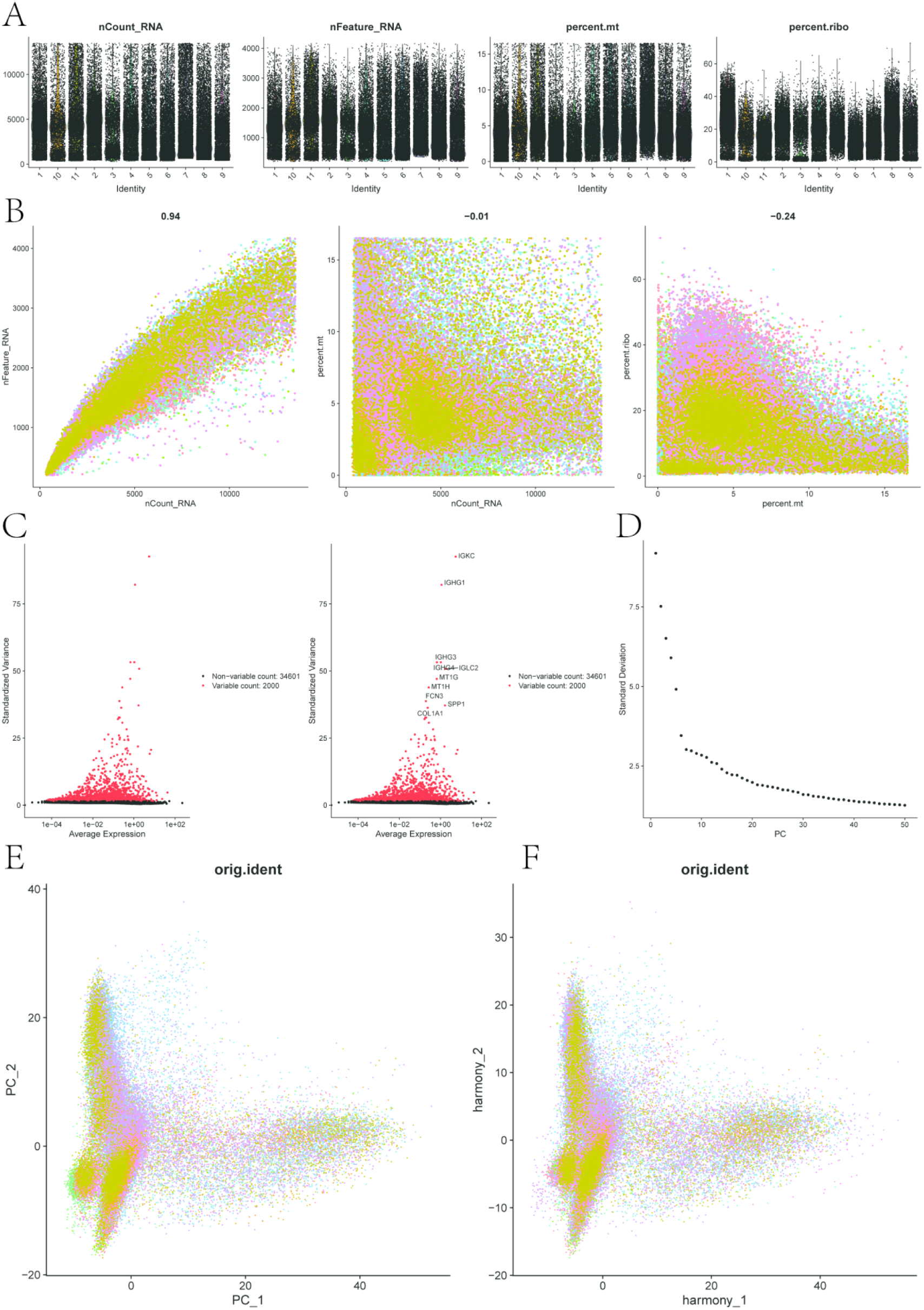
Single-cell preprocessing (A) Single-cell quality control showing the number of cells, number of genes, and sequencing depth for each sample. (B) The left panel shows the relationship between cellular sequencing depth and mitochondrial content, and the right panel shows the relationship between sequencing depth and number of genes, both of which are positively correlated. (C) We identified genes with significant differences among cells and plotted the feature variance graph. (D) Variance ranking plot for each PC. (E-F) PCA visualization and PC distribution, where points represent cells and colors represent samples.

### 2.9 Expression of NABP2 in cell subgroups

In this study, each subgroup was further annotated, and 12 clusters were annotated into 10 cell categories: CD8+ T cells, NK cells, Macrophages, Hepatocytes, Neutrophils, Endothelial cells, Monocytes, CD4+ T cells, B cells, and Fibroblasts (Figure 10A). Additionally, we presented bubble plots of the classical markers of these 10 cell types (Figure 10B) and bar charts of cell proportions (Figure 10D). Furthermore, we analyzed the expression of NABP2 in the 10 cell types at the single-cell level (Figures 11A-C) and displayed the proportions of each cell type in NABP2 positive and negative groups (Figure 11D).

**Figure 10.**
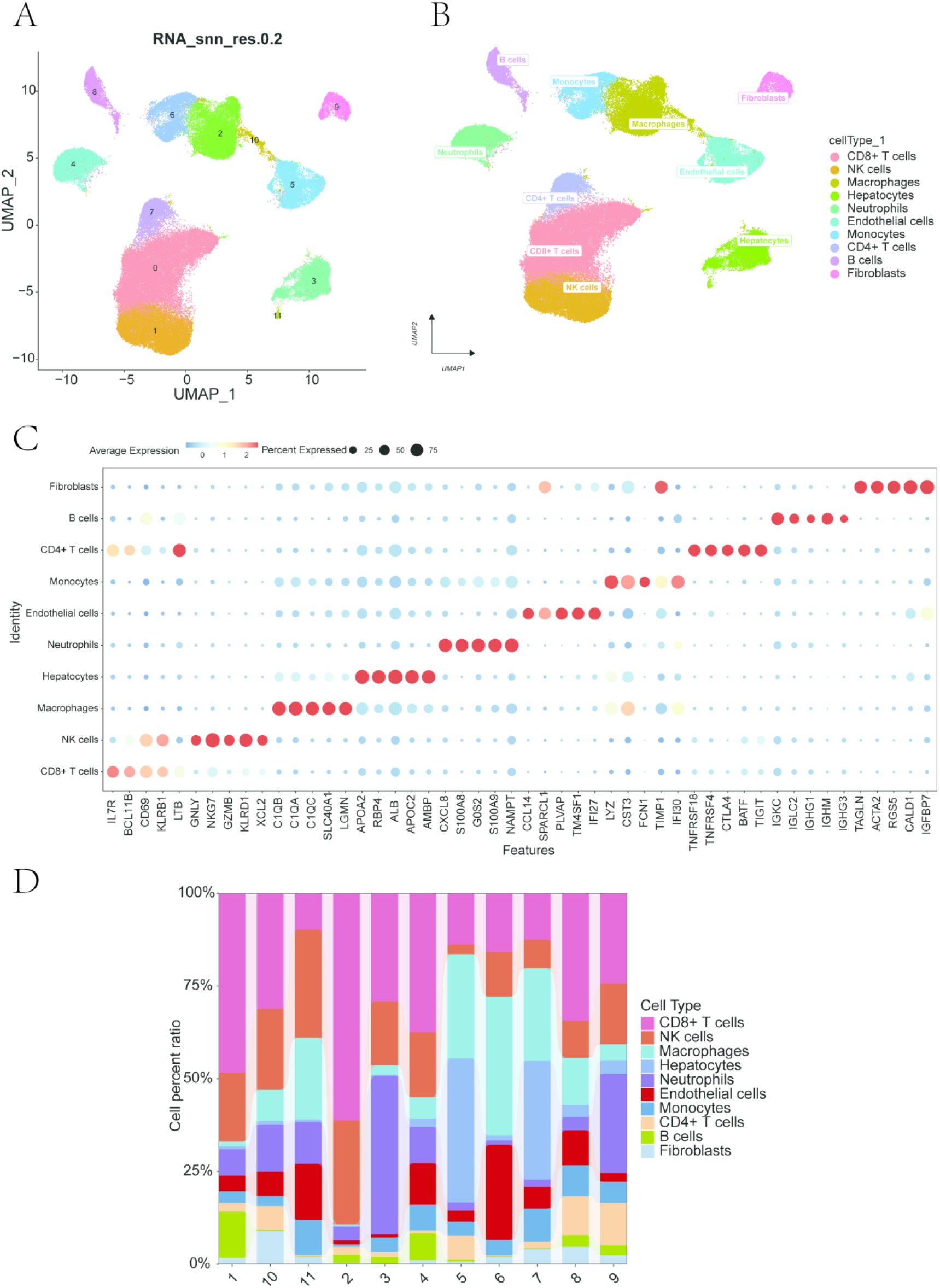
cell annotation results (A) Based on the important components available in PCA, we divided the cells into 12 clusters using the UMAP algorithm. (B) cell annotation results of the 12 clusters, which were annotated into 10 cell types, namely CD8+ T cells, NK cells, Macrophages, Hepatocytes, Neutrophils, Endothelial cells, Monocytes, CD4+ T cells, B cells, and Fibroblasts. (C) Bubble plot of Doplot showing 10 cell types and their cell markers. (D) Differences in the proportion of 10 cell types among 11 samples.

**Figure 11.**
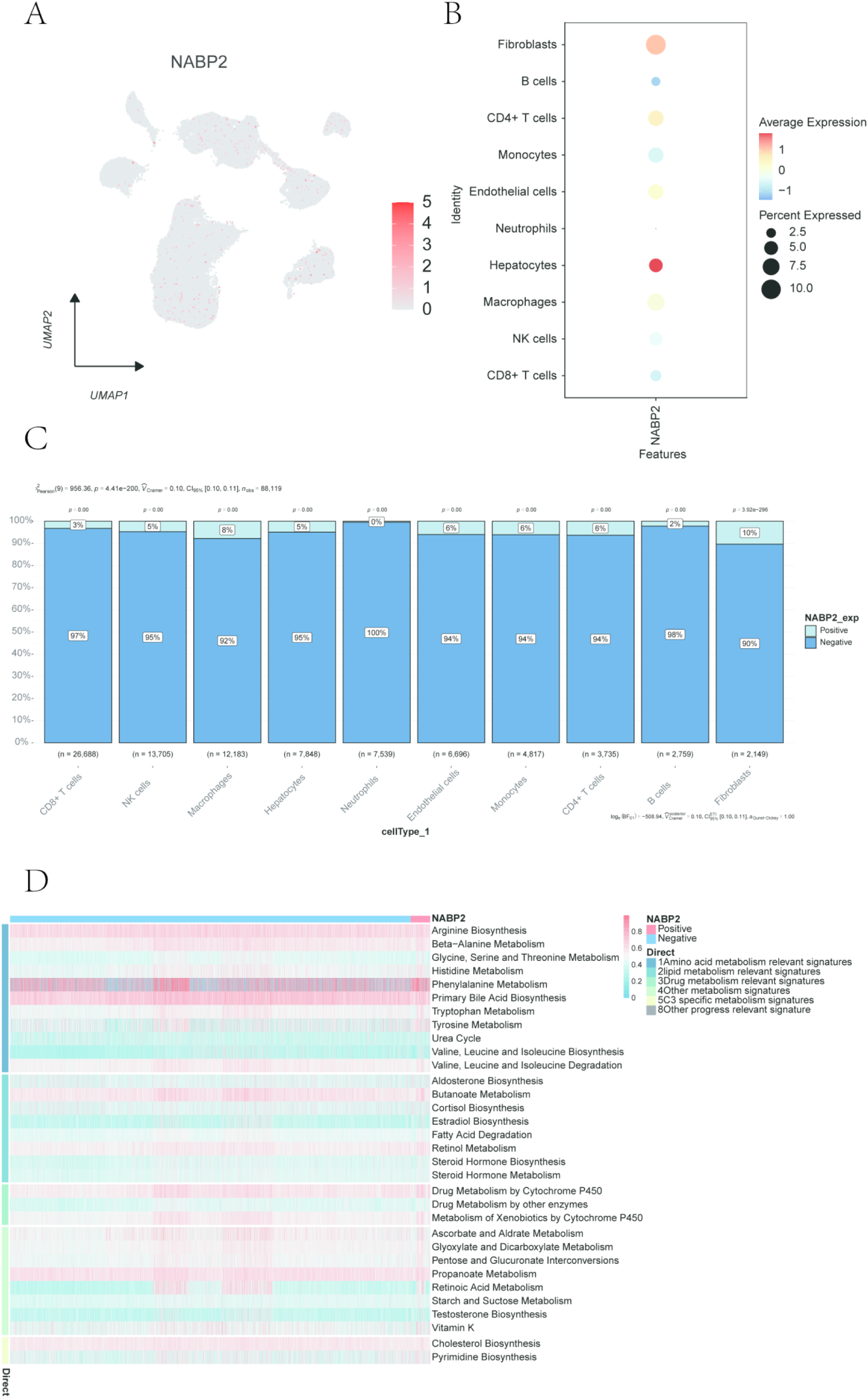
Single-cell expression profiles and correlation with metabolic pathways (A) Scatter plot of the expression profiles of key genes in single cells. (B) Bubble plot of the expression profiles of key genes in single cells. Blue represents low expression, and red represents high expression. (C) Differences in the proportion of each cell type between NABP2 positive and negative groups. (D) Correlation between NABP2 gene and metabolic pathways, with blue indicating low expression and red indicating high expression.

### 2.10 Interaction between NABP2 and cell subgroups

We analyzed the ligand-receptor relationships of the features in NABP2 positive and negative groups from the single-cell expression profiles of liver cancer using the cell Chat software package, respectively. In positive cells, we found complex interaction pairs among these cell subgroups (Figure 12A). In addition, our statistics showed that cells such as Macrophages had closer potential interactions with other cells. Therefore, the ligand-receptor status during the interaction between Macrophages and various cell subgroups in NABP2 positive cells (p<0.01) is shown (Figure 12B). In negative cells, we found complex interaction pairs among these cell subgroups (Figure 12D). In addition, our statistics showed that cells such as Macrophages had closer potential interactions with other cells. Therefore, the ligand-receptor status during the interaction between macrophages and various cell subgroups in NABP2-negative cells (p<0.01) is shown (Figure 12E).

**Figure 12.**
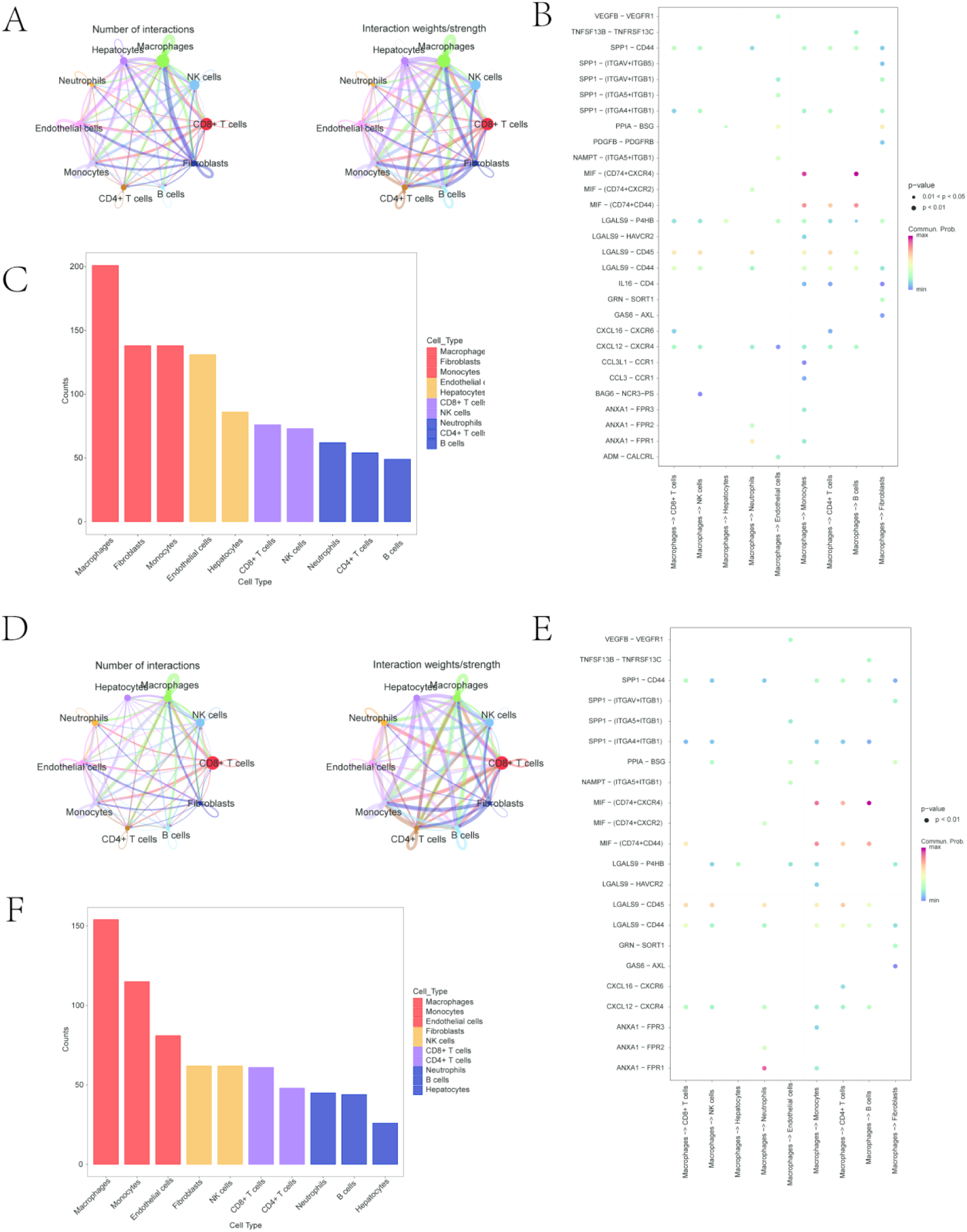
cell Communication (A) cell interaction network among 10 types of positive cells, where the edge width indicates the probability and strength of communication between cells. (B) Comparison of the total number of interactions in the communication network among 10 types of positive cells, decreasing from left to right, with Macrophages being the strongest. (C) Bubble plot of receptor-ligand between positive cells and cells. (D) cell interaction network among 10 types of negative cells, where the edge width indicates the probability and strength of communication between cells. (E) Comparison of the total number of interactions in the communication network among 10 types of negative cells, decreasing from left to right, with Macrophages being the strongest. (F) Bubble plot of receptor-ligand between negative cells and cells.

### 2.11 Correlation between NABP2 and metabolic pathways

In this study, the enrichment scores of different metabolic pathways in each cell were calculated by the software package GSVA. Then, according to the expression grouping of NABP2, the results showed that the positive group of the NABP2 gene had higher activities in pathways such as Beta-Alanine Metabolism and Butanoate Metabolism. (Figure 12C, F)

## 3 Discussion

China has the highest incidence rate of liver cancer globally. In 2020, Chinese liver cancer patients accounted for 45.3% of the world’s total liver cancer patients, and liver cancer deaths accounted for 47.1% of the global total. This severe reality indicates a higher demand for treatment and nursing resources among liver cancer patients [24]. Currently, the 5-year survival rate of non-surgical treatment is 20%, and the success rate of treatment using special methods is not particularly high. Finding a new type of treatment is necessary for the medical industry. Through our research, it was found that NABP2 and its related metabolic pathways may provide a new direction for the treatment of liver cancer.

As one of the emerging fields in oncology research in recent years, the tumor immune microenvironment is of great significance for in-depth study of the pathogenesis and metastasis pathways of tumors [25]. The tumor immune microenvironment mainly consists of immune cells in tumor tissues and the cytokines they secrete, such as regulatory T cells, which have immunosuppressive properties and can restore immune homeostasis after inflammation by inhibiting the activity of various immune cells (CD8+ and CD4+ effector T cells, NK cells, DC cells) [26]. In addition, the tumor immune microenvironment also has effects such as inhibiting tumor immune responses and affecting the adhesion and migration of tumor cells [27]. NABP2 affects the growth and development of lymphocytes, dendritic cells, and macrophages in the immune system [28]. NABP2It can bind and interact with receptors on immune cells, affecting cell activation and immune responses. Through our research, we found that NABP2 is significantly positively correlated with cells such as T cells regulatory (Tregs) and Macrophages M0, and significantly negatively correlated with cells such as Monocytes and B cells naïve.

Through various bioinformatics analyses, we found that NABP2 affects the occurrence and development of liver cancer and its anti-tumor immune microenvironment [29]. Targeted drugs such as sorafenib and regorafenib can specifically target certain molecular characteristics of tumor cells, resulting in inhibitory effects on tumor cell growth. Platinum-based drugs such as cisplatin and carboplatin destroy the structure and function of DNA based on the principle of binding to DNA and causing interstrand or interstrand cross-linking [30]. Since NABP2 is involved in DNA metabolism, these drugs may indirectly affect the function of NABP2 and exert anti-cancer effects. The above-mentioned drugs still inevitably have problems of inaccurate targeting and high cost. Therefore, we further explored the correlation between the NABP2 gene and the sensitivity to chemotherapy drugs to screen chemotherapeutic drugs.

To investigate the molecular regulatory mechanisms and pathways by which the NABP2 gene is involved in liver cancer, we performed GSVA and GSEA analyses respectively. The GSVA results showed that the NABP2 gene may affect the disease progression of liver cancer by participating in signaling pathways such as E2F-TARGETS, DNA-REPAIR, and MYC-TARGETS-V1. Among them, the E2F-TARGETS signaling pathway is a signal transduction pathway closely related to cell cycle regulation and plays a key role in the occurrence and development of various cancers. E2F family transcription factors can activate or inhibit the expression of a series of cell cycle-related genes, including those involved in DNA synthesis, S-phase progression, and mitosis. The E2F family participates in most human malignant tumors by activating the transcription of cell cycle-related genes.E2F has multiple influence pathways in liver cancer. For example, it can induce macrophage infiltration by promoting the overexpression of certain cancer cell targets such as MTCH1 and POLA2 [31**] [**32], leading to poor prognosis of liver cancer; E2F can also promote the proliferation of liver cancer cells by promoting the transcription of related genes [33]. Based on the enrichment analysis, NABP2 is involved in E2F targets, so we believe that NABP2 may have certain functions in liver cancer treatment. DNA-REPAIR is a signaling pathway related to DNA repair, which can recognize and repair damaged DNA molecules. The function of this signaling pathway in maintaining genome stability plays a certain role in cancer treatment. MYC-TARGETS-V1 is a signal transduction pathway closely related to cell growth, proliferation and metabolism. Studies have shown that in cancer-related research, gene amplification and overexpression of MYC often occur. In addition, the MYC signaling pathway can also re-program cell metabolism to enable tumor cells to adapt to the needs of rapid growth and division, thereby accelerating cancer progression.GSEA results indicated that the NABP2 gene can also be enriched in signaling pathways such as DNA replication, Proteasome, and Spliceosome, suggesting that this gene plays a key role in a series of molecular regulatory processes including DNA replication, protein degradation, and mRNA splicing. Therefore, we have demonstrated that the NABP2 gene is involved in numerous cancer-related signaling pathways; however, the specific mechanism of its involvement remains unclear and requires further in-depth investigation. Future research needs to deeply analyze the downstream effector molecules and signal networks of NABP2 by combining experimental methods. These limitations do not affect the reliability of the core conclusions of this study, but rather point out the direction for future research.

According to this study, we can conclude that NABP2 drives the malignant progression of liver cancer and reshapes the immunosuppressive microenvironment through the E2F/MYC axis. This article is the first to reveal that NABP2 simultaneously possesses the functions of protecting the genome and coordinating immunometabolism in liver cancer. GSVA/GSEA confirmed that high NABP2 expression enriches E2F targets and MYC signaling. In single-cell validation, the expression of E2F target genes was increased by 32-fold (p=1.4e-8) and was positively correlated with the proportion of G2/M phase cells (r=0.67), suggesting that NABP2 provides protection for E2F/MYC-driven cell cycle dysregulation by maintaining genomic stability. Meanwhile, TCGA analysis showed a significant increase in Tregs and M2 macrophage infiltration in the high NABP2 expression group, and cellChat analysis revealed that NABP2 macrophages highly express TGFB1 and IL10, which interact with TGFBR2/IL10RA on Tregs to activate the immunosuppressive network. This group is absent in the NABP2-group, indicating that NABP2 directly regulates immunosuppressive cell crosstalk.

Peng [29] et al.’s study supports the role of NABP2 as an oncogene in liver cancer and suggests its promotion of metastasis. The present study not only validates its cancer-promoting effect but, more importantly, expands the understanding of its functions, particularly its roles in regulating the immune microenvironment of liver cancer (promoting Tregs and M2 macrophage infiltration) and metabolic pathways, which provide a more comprehensive explanation for the discovery of NABP2’s cancer-promoting mechanisms. However, the specific molecular mechanisms by which NABP2 regulates immune cell infiltration and metabolism, such as whether it directly affects immune cell functions or indirectly acts by altering tumor cell secretory factors, remain to be further explored.

Of greater note in this study, we also specifically focused on the correlation between NABP2 and metabolic pathways. Similarly, through GSVA analysis, the results showed that the NABP2 positive group had higher activity in pathways such as Beta-Alanine Metabolism and Butanoate Metabolism. Therefore, from the perspective of metabolomics, we can further explore the specific regulatory mechanisms and differential expression of the NABP2 gene in β-alanine and butyrate metabolism, and also further investigate the specific roles of these two metabolic pathways in the occurrence and development of liver cancer, which may be a very promising therapeutic option. Despite some limitations, this study systematically depicts the multifunctional role of NABP2 in liver cancer, linking it to DNA metabolism, shaping of the immunosuppressive microenvironment, andmetabolic pathway activation. NABP2 or its regulatory pathways are expected to become new targets for liver cancer treatment, providing a theoretical basis for exploring strategies targeting NABP2 in combination with immune checkpoint inhibitors or chemotherapy.

